# Coarse-scale Optoretinography(CoORG) with extended field-of-view for normative characterization

**DOI:** 10.1101/2022.09.08.507165

**Authors:** Xiaoyun Jiang, Teng Liu, Vimal Prabhu Pandiyan, Emily Slezak, Ramkumar Sabesan

## Abstract

Optoretinography (ORG) has the potential to be an effective biomarker for light-evoked retinal activity owing to its sensitive, objective, and precise localization of retinal function and dysfunction. Many ORG implementations have used adaptive optics (AO) to localize activity on a cellular scale. However, the use of AO restricts field-of-view (FOV) to the isoplanatic angle, necessitating the montaging of multiple regions-of-interest to cover an extended field. In addition, subjects with lens opacities, increased eye movements and decreased mobility pose challenges for effective AO operation. Here, we developed a coarse-scale ORG (CoORG) system without AO, which accommodates FOVs up to 5.5 deg. in a single acquisition. The system is based on a line-scan spectral domain OCT with volume rates of up to 32 Hz (16,000 B-frames per second). For acquiring ORGs, 5.5 deg. wide OCT volumes were recorded after dark adaptation and two different stimulus bleaches. The stimulus-evoked optical phase change was calculated from the reflections encasing the cone outer segments and its variation was assessed vs. eccentricity in 12 healthy subjects. The general behavior of ΔOPL vs. time mimicked published reports. High trial-to-trial repeatability was observed across subjects and with eccentricity. Comparison of ORG between CoORG and AO-OCT based ORG at 1.5°, 2.5°, and 3.5° eccentricity showed an excellent agreement in the same 2 subjects. The amplitude of the ORG response decreased with increasing eccentricity. The variation of ORG characteristics between subjects and versus eccentricity was well explained by the photon density of the stimulus on the retina and the outer segment length. Overall, the high repeatability and rapid acquisition over an extended field enabled the normative characterization of the cone ORG response in healthy eyes, and provides a promising avenue for translating ORG for widespread clinical application.

## 1. Introduction

Optoretinography (ORG), a term first introduced in the early 1980s [1], describes the non-invasive, optical imaging of light-induced functional activity in the retina and stands to be a precise diagnostic biomarker for retinal disease. Compared to other ophthalmic exams, ORG has significant advantages in providing a non-invasive, functional, objective, and sensitive assay of retinal function in response to light stimuli. Recent work has employed a variety of optical imaging modalities for acquiring the ORG, including SLOs[2-4], fundus cameras[5, 6], and OCT[7-18]. These can be broadly categorized as those imaging the light-evoked backscattered intensity[2-6, 8-11], and those analyzing the backscattered optical phase [7, 12-14, 17, 18].

The photoreceptors have been the focus of prior work on ORGs with a couple of exceptions[19, 20]. Cone photoreceptors are readily imaged with adaptive optics (AO). When combined with the aforementioned imaging modalities of SLO, fundus cameras and OCT, AO provides cellular-scale access to both the structure and function of individual photoreceptors. Accordingly, a majority of previous ORG implementations have employed AO, enabling for instance, the efficient classification of cone sub-types, and sensitive assessment of cellular scale dysfunction in retinitis pigmentosa[13-15, 17, 21].

However, the use of AO poses some challenges that restrict applicability and throughput for ORG. Typically, wavefront measurement and correction is performed for one field point when the wavefront beacon is not scanned, or the average of field points across the imaging field for a scanning system. The eye‘s aberrations change across the visual field. Field angles beyond the isoplanatic angle of the eye‘s optics at any eccentricity remain sub-optimally corrected. This restricts the overall field-of-view (FOV) imaged and necessitates the montaging of multiple regions-of-interest to improve retinal coverage. In addition, subjects with lens opacities, increased eye movements and decreased mobility pose challenges for effective closed-loop AO operation.

Regardless, AO incorporated into SLO and OCT has allowed characterizing the fundamental features of the cone ORG response. Using an AOSLO, Cooper et al.[3] showed that the action spectrum of light-evoked changes in backscattered intensity in cones follows the photopic luminosity function. In phase-resolved OCT, few features of the cone outer segment optical path length change (ΔOPL) in response to stimuli have been noted[12, 14, 17]. It is known that cone outer segments exhibit a rapid shrinkage in ΔOPL that is followed by a slow expansion. The magnitude and slope of this light-evoked signal increases with stimulus flux. For these features to find applicability in the study of retinal diseases, or to understand their physiological mechanisms, it is critical to assess the degree to which they vary within and between individuals. To that end, it is imperative to perform a normative characterization of the ORG.

Here, we report on a coarse-scale ORG (CoORG) system without AO and demonstrate its viability for ORG recording of FOV up to 5 degrees in a single acquisition. We assess how the CoORG compares with the high-resolution AO ORG measurements, how the ORG varies versus eccentricity, photon flux and between individuals. This work thus initiates the normative characterization of phase-based cone ORGs, and will aid in establishing baselines against which diseases and therapeutics can be compared.

## 2. Methods

### 2.1 System layout and specifications

The CoORG system is based on the first generation lens-based adaptive-optics-(AO) line-scan spectral domain OCT described previously [13], with key differences optimized at expanding the imaging field. First, it does not include an AO module − the deformable mirror and wavefront sensor. To accommodate an extended field, it was designed to have a smaller diameter entrance pupil at the eye equal to 3 mm. Larger 2-inch diameter achromatic lens-based telescopes were used to relay pupil and imaging planes, to avoid beam vignetting. In the detection channel, an anamorphic telescope was used to optimize light collection, lateral and spectral resolution in the AO system previously. Here, such an anamorphic optimization was not needed given the preference to maintain a larger field over cellular resolution.

Figure 1 shows the system layout. An 840±25nm super luminescent diode (MS-840-B-I-20, Superlum, Ireland) was collimated by a reflective fiber collimator to form an 8.5mm diameter beam. An achromatic cylindrical lens (CL1) was used to form a line at the entrance pupil (*P1*). A 30/70 (reflectance/transmission) beam splitter (BS) was used to split the beam into the sample and reference arm. The sample arm beam was relayed by three pairs of afocal telescopes (L1 to L6) to the eye‘s pupil. The entrance pupil (*P1*), galvo-scanner (*P2*), an extra pupil plane for refractive error correction (*P3*) and the eye‘s pupil (*P4*) were optically conjugated using these achromatic lens-based telescopes. L1 to L6 were slightly tilted to remove the back reflection from each lens and minimize the aberrations so introduced in the system due to the tilt. A camera placed at *R*5 was used to monitor the back reflected point-spread-function from a model eye, and the lenses were serially tilted in order to minimize the PSF extent. Trial lenses placed at *P*3 were used to correct residual refractive error of the system and the subject‘s eye. The reference arm beam was re-collimated by the second achromatic cylindrical lens (CL2), then passed through two afocal telescopes (L7 to L10) to reduce the diffraction due to the long-distance propagation. A glass window (DG) was inserted in the reference arm to compensate the difference in dispersion between the sample and reference arms. A mirror M9 was placed on a translation stage and adjusted to match path lengths between the sample and reference arms.

**Fig. 1.**
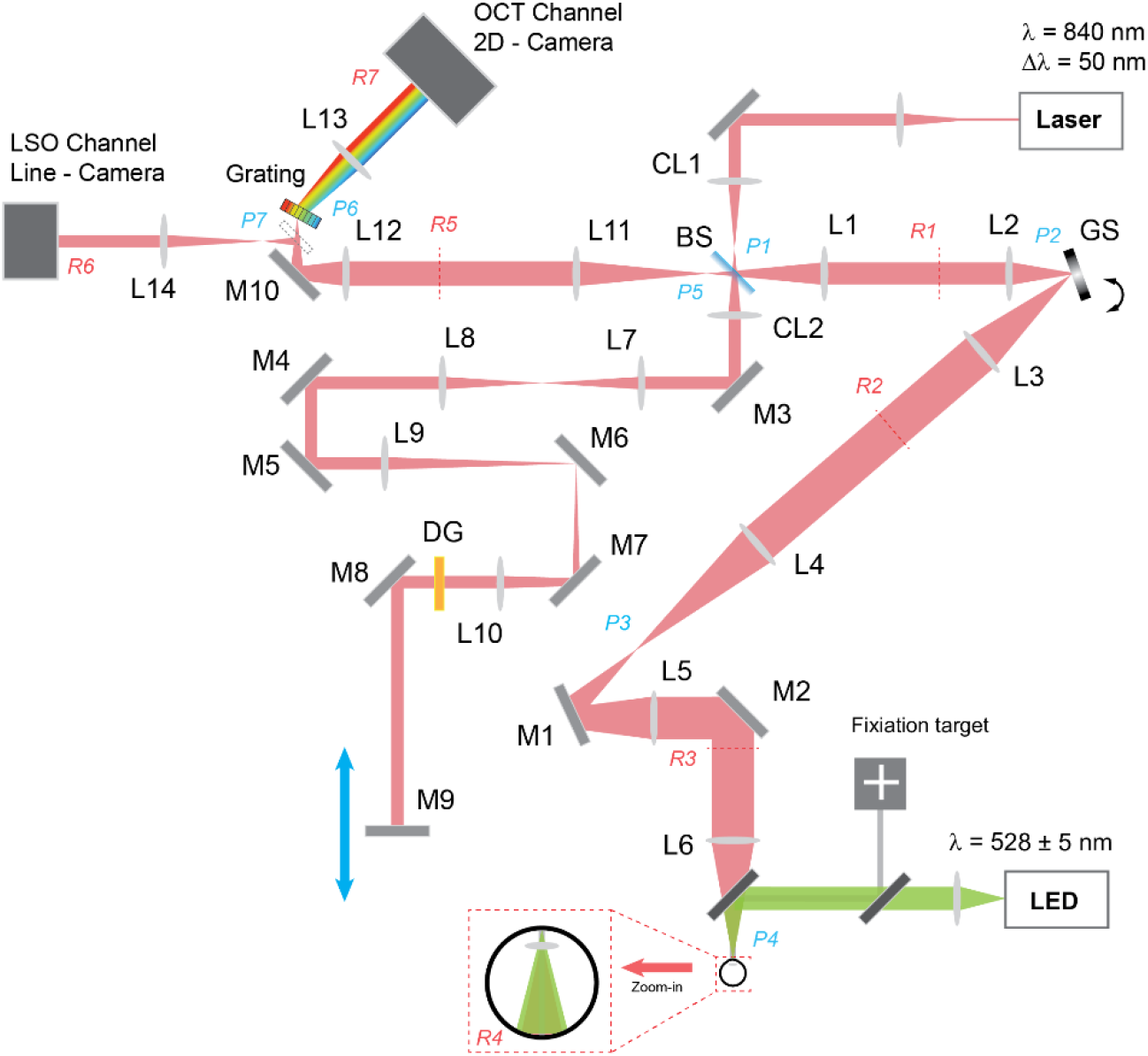
CoORG system layout. The layout is not drawn to scale, and the beam follows the general path on the optical table. The beam is shown along one dimension in illumination where the linear field is focused on the pupil planes. The orthogonal view is not shown. The beam path in detection corresponds to the case where a mirror is placed at the eye‘s pupil. L: spherical lens, CL: cylindrical lens, M: mirror, GS: galvo scanner, BS: beam splitter, DG: dispersion compensation glass window, LSO: line-scan ophthalmoscope, OCT: optical coherence tomography, P: pupil plane, R: retinal plane, LED: light-emitting diode.

The sample and reference arms were combined in the detection arm. An iris was placed at the pupil plane (*P5*) and adjusted to correspond to a 3mm beam diameter at the eye‘s pupil plane (*P4*). A slit was added at the imaging plane (*R5*) to minimize remaining back reflections and stray light reaching the detectors and to provide partial confocality. An interchangeable mirror was used to switch between the line-scan ophthalmoscope (LSO) and the line-scan OCT modalities. A 1200 line-pairs/mm diffraction grating was placed at pupil plane *P*6 to disperse the interference signal into its spectral components on the OCT detector. A fast 2D CMOS camera (Photron, FASTCAM NOVA S6) was used to capture the OCT signal (*R*7) and a line camera (Basler, sprint, spL2048-70 km) was used to capture the LSO signal (*R*6). An LED (528 ± 5nm) was introduced into the eye in Maxwellian view to serve as the light stimulus for ORG experiments and combined with the imaging light using a long pass filter. The system specifications are listed in Table.1

**Table 1.** System specification.

|  |  |
| --- | --- |
| Lateral resolution | 3.6 $\mu\text{m}$ |
| Axial resolution in air | 6.23 $\mu\text{m}$ |
| Axial depth in air | 2 mm |
| FOV | 5.5 deg |

### 2.2 Imaging protocol

Twelve subjects free of retinal disease (ages 20-42 years old) were recruited for the study after an informed consent. The research was approved by the University of Washington institutional review board and conducted according to the tenets of the Declaration of Helsinki. The pupil was dilated using Tropicamide 1% ophthalmic solution (Akorn Inc.). Without dilation, pupil miosis due to the bright stimulus light impairs comparative analysis between subjects. Instead, a maximally dilated pupil was used for all measurements, and its diameter (along with axial length) was measured to incorporate into the stimulus photon density calculation for each subject (section 2.4).

A schematic of the experimental procedure is outlined in Figure 2. First, the axial length of each subject was measured using Zeiss IOLMaster (Carl Zeiss Meditec AG, Germany) while waiting for pupil dilation. Next, a 15° × 5° FOV OCT image stack was taken along with a 30 ° ×30 ° SLO image (Fig.2B) using a commercial OCT system (Heidelberg OCT Spectralis, Heidelberg Germany). The conventional OCT & SLO images were used as a preliminary guide to locate the region where the CoORG data were acquired (Fig.2D). The subject was then seated in the CoORG system and aligned to the system‘s exit pupil. Pupil size was measured from the pupil image acquired when the subject‘s eye was well focused at the exit pupil of the imaging system (Fig.2C). A live real-time LSO image stream was used to locate the region of interest (ROI) and find best focus of the retina by adding trial lenses at the pupil plane *P3* (seen in Fig.1). Then, the system was switched to line-scan OCT mode and ORGs were recorded as described below. The OCT *en face* image at the inner-outer segment junction (ISOS) was registered with the clinical SLO image. Using the foveal pit as the reference for zero-degree eccentricity, the ORG signals were extracted from annular regions-of-interest surrounding the fovea, in the range from 0.5 deg to the edge of the imaging field, typically 4.5 − 5 deg. While the FOV of the instrument is 5.5 deg., eye movements slightly decrease the field available to reliably average over to obtain ORGs. The details of the data processing are described in section 2.3.

**Fig. 2.**
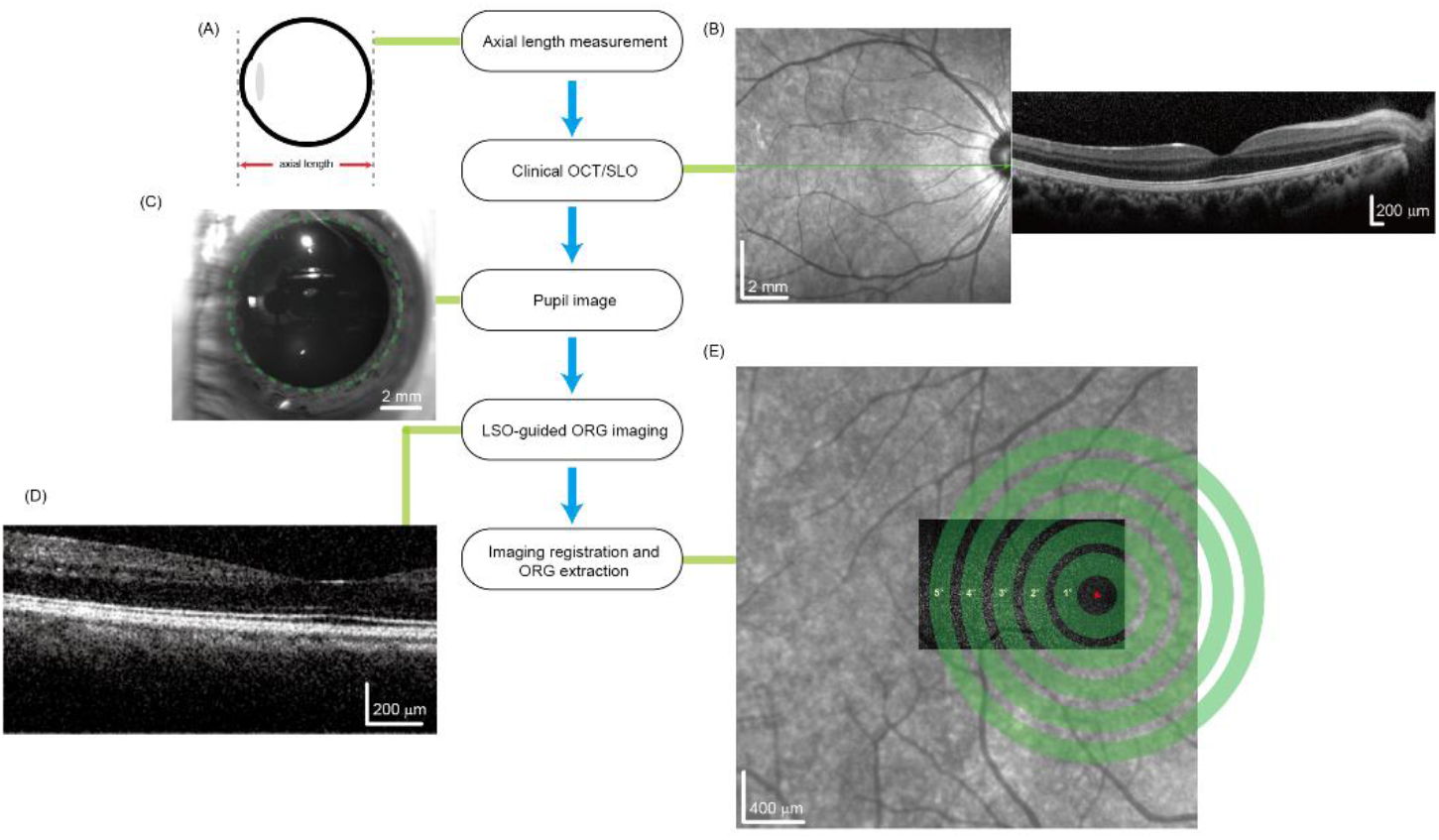
Experimental flow. (A) Axial length was measured by Zeiss IOLMaster (B) Clinical SLO and OCT was taken for registration and finding the foveola; (C) Pupil image was taken for measuring the pupil size; (D) CoORG system B scan example. (E) ORG extraction region (green ring) superimposed on the registered IS/OS *en face* image and clinical SLO image. Red cross indicated the center of the fovea.

For acquiring ORGs, the subject was dark adapted for one minute. Then, 50 OCT volumes were recorded at a rate of 30 Hz. The LED stimulus light was delivered at the 11^th^ volume and lasted for either 10 or 20ms for low and high bleach levels respectively. Five repeat trials were taken for each bleach level. Each trial including the time for dark adaptation took about 2 minutes. In all, the ORG imaging session lasted ∼20 minutes for each subject including both bleaches. One subject‘s right eye was imaged with fixation at 2.5 deg temporal and 2.5 deg inferior to check for meridional dependence. The rest of the subjects were imaged with fixation at 2.5 deg temporal eccentricity.

It was important to compare the CoORG against measurements taken in individual cones with AO. For this, a reflective line-scan AO-OCT described previously was used[15]. Two subjects were imaged with AO at 1.5°, 2.5° and 3.5° inferior eccentricity following the same ORG protocol with the higher of two bleach levels.

### 2.3 Data processing

The OCT data processing followed steps described in Ref. [13]. Briefly, typical OCT reconstruction − k-space remapping and fast Fourier transform − was conducted to get B-scan stacks for each volume in a given trial. An open-source segmentation software [22] was used to identify the outer retinal layers, adapted and optimized for the line-scan OCT. The five repeat trials were co-registered together based on a strip-based image registration algorithm [23] on the basis of the enface ISOS layer image. *En face* images at the ISOS and cone outer segment tips (COST) were extracted from the registered volume and manually inspected to confirm accurate segmentation and registration. The center of the fovea (red mark in Fig.2E) was determined in the clinical SLO and *en face* ISOS image based on the corresponding OCT B-scan foveal pit location. The dependence of the ORG vs. eccentricity was computed from this foveal position as the zero-degree eccentricity reference. The ORG signal was averaged in each 0.5° annular ring, with a step size in eccentricity of 0.5°. For illustration, Fig.2E shows five rings corresponding to integer steps in eccentricity. The steps for ORG processing followed previous work[13], based on stimulus induced phase difference analysis between the ISOS and COST. For measurements with AO, the mean of individual cones was obtained and compared against averaged ORG responses from CoORG at the same eccentricity.

### 2.4 Calculating photon density at the retina

The ORG signal characteristics depend acutely on the stimulus photon density on the retina. To capture any inter-individual differences, an individualized calculation of photon density was performed. First, the spectrum and power of the LED were measured using a spectrometer and power meter, respectively. The angular subtends of the stimulus field was calculated by measuring the ratio of its absolute physical size a given distance away from the entrance pupil. These three measured parameters remained constant for each subject. The LED beam size at the entrance pupil was greater than the largest dilated pupil; hence the power was scaled by each subject‘s measured pupil area (Fig.2C). The measured power in units of Watts was integrated across the spectrum and converted to photons.s^-1^ using the physical constants for speed of light in vacuum, c = 3.0e8 m/s and the Planck‘s constant h = 6.626e-34 J.s. Absorption by the lens[24] and macular pigment[25] were accounted for using published values to estimate the quanta per second reaching the retinal surface. Dividing this value by the stimulus area in angular units, provided an estimate of photons.sec^-1^.deg^-2^. The LED pulse width (10 or 20 ms) and the axial length of the subject were used to convert these units to photons.µm ^-2^ assuming a scaling factor of 291 µm per degree for a 24 mm axial length.

## 3. Results

### 3.1 OCT cross-sectional images

Figure 3A shows an example of a B-scan taken by CoORG system extending from the fovea up to 4.5° temporal eccentricity. Figure 3B shows the intensity profile of A-lines at different eccentricities showing the different peaks corresponding to the typical outer retinal layers. From the fovea to 1.5°, there are three dominant peaks, corresponding to the ISOS, COST and retinal pigment epithelium (RPE). Starting from 2° eccentricity, a fourth peak appears between the COST and RPE, corresponding to the tips of the rod outer segments (ROST). This demonstrates the ability of the instrument to capture B-scans with high fidelity showing the distribution of the major retinal layers.

**Fig. 3.**
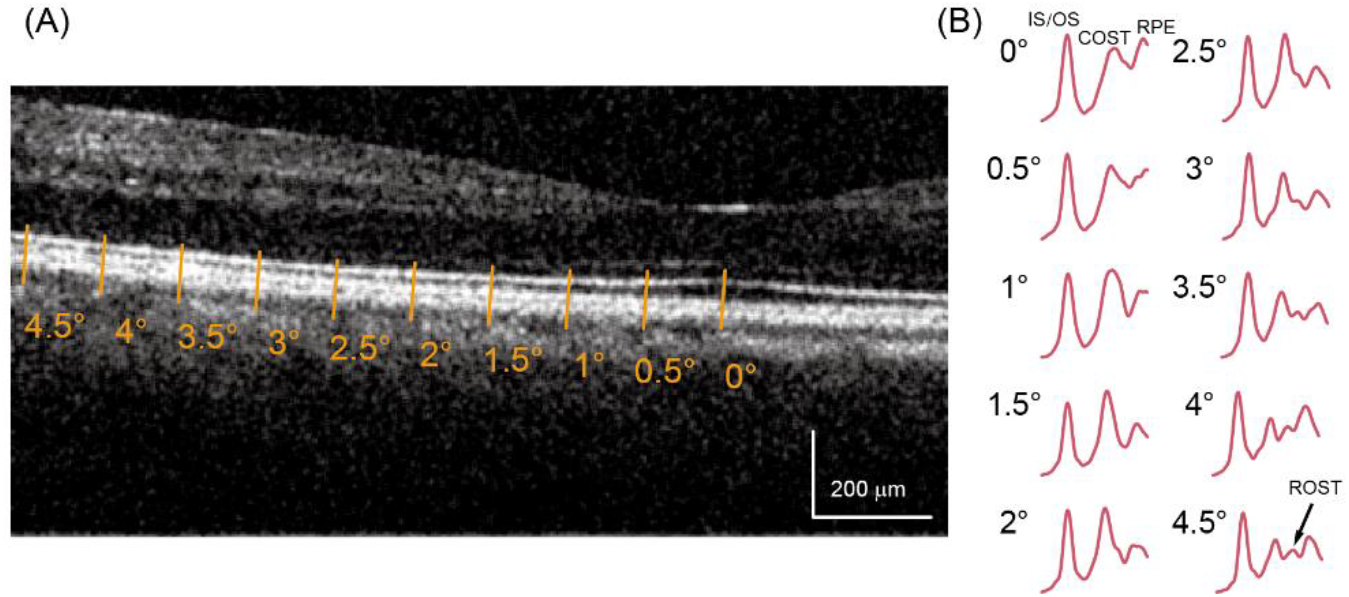
Example of a B-scan acquired from the CoORG instrument. Eccentricities are indicated on the B-scan, and the corresponding intensity profiles along the A-scan are shown in (B). The peaks correspond to the major retinal layers(details in text).

### 3.2 ORG trial-to-trial repeatability

The repeatability was assessed across 5 repeated measurements and two bleach conditions summarized in Fig. 4 for a subject. Each panel shows the change in ΔOPL in the outer segment on the y-axis, and the time after stimulus onset on the x-axis. The general trend of the curves follows published reports[12, 14, 17], where immediately after light onset, the ΔOPL rises rapidly and plateaus in about 300ms. The saturated amplitude and the slope of rise of ΔOPL are known to increase with stimulus bleach level[14] and these characteristics are observed here as well for each eccentricity. In addition, it is noted that with increasing eccentricity, the saturated amplitude and slope exhibit a reduction in their magnitudes. Figures 4A and B show the high and low bleach levels from 1° − 4° temporal eccentricity. The different traces in each panel correspond to a single measurement. Each panel in Fig.4A and B demonstrates high trial-to-trial repeatability. The mean across all 5 trials for both bleach levels are shown in Fig.4C and D, respectively. The time-averaged standard deviations across trials were calculated for each eccentricity in 0.5° steps and ranged between 6.5 to 22.2 nm for the high bleach and between 4.8 and 13.0 nm for the low bleach. The average standard deviation vs. eccentricity was ±13.34 nm and ±9.68 nm for high and low bleach level, respectively in this subject. The same was calculated for all subjects, and the average standard deviation between subjects was ±8.3 nm (range: 2.1 - 23.5nm) and ±4.5nm (range :1.0-18.8nm) for 20 and 10ms stimulus flashes respectively. Overall, this indicates that the ΔOPL vs. time was highly repeatable in CoORG and follows the previously reported measurements closely.

**Fig. 4.**
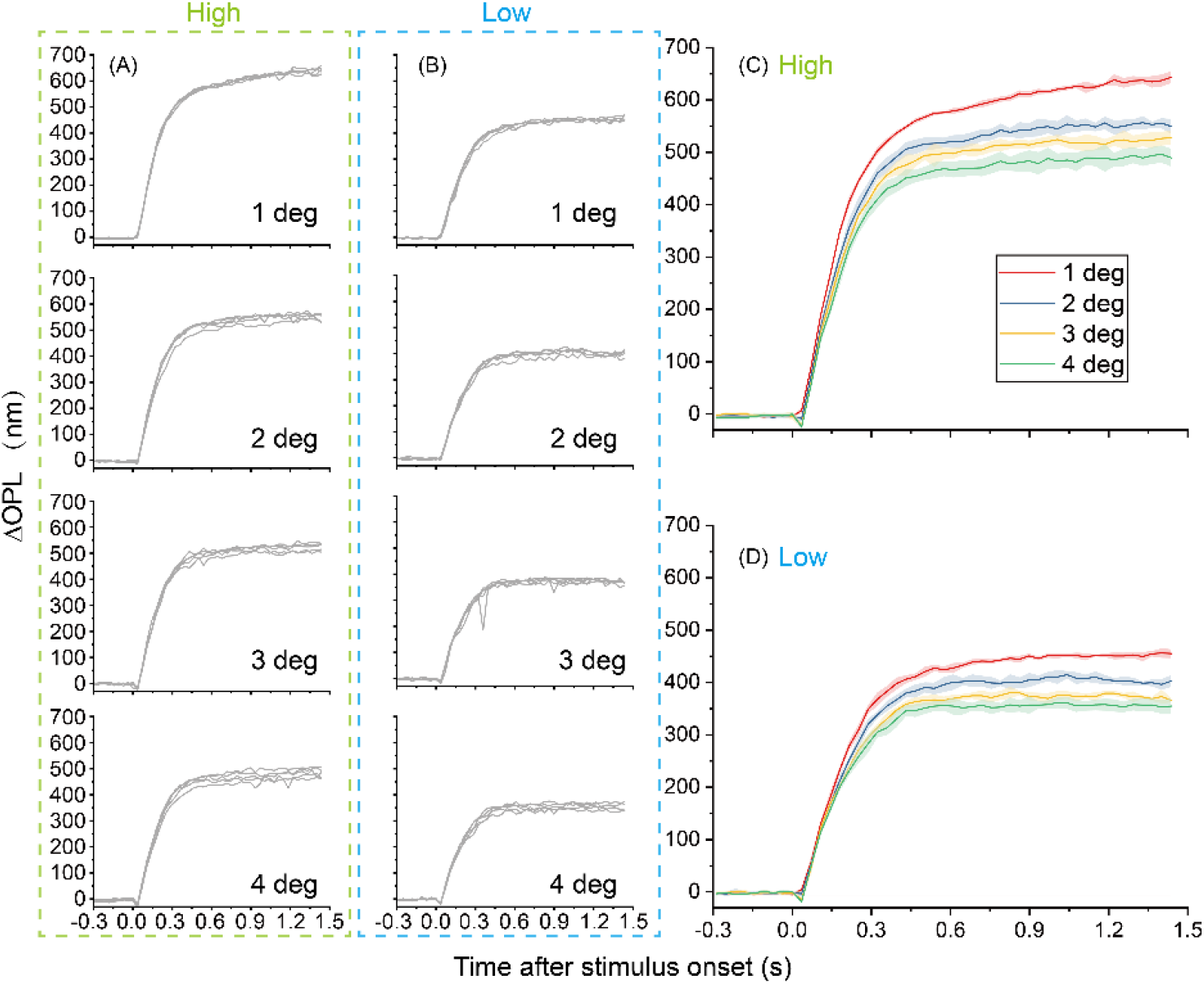
Light-evoked ΔOPL for a subject along the temporal meridian. Column A and column B show plots of 5 successive repeat trials at different eccentricities for high and low bleach levels, respectively where each curve represents a single trial. The high and low bleach correspond to 8.8×10^6^ and 4.4×10^6^ photons/µm^2^ respectively for this subject. Plot C and D are the mean distributions of the five repeat measurements for both bleach levels at different eccentricity. The shaded area surrounding the solid traces represent ±1 standard deviation.

### 3.3 Dependence of ORG on retinal meridian

To assess any potential dependence on retinal eccentricities at different meridians, the same subject was imaged along the inferior and temporal eccentricity, at the same high bleach level. The maximum saturated ΔOPL was calculated from the ORG trace by taking the mean ΔOPL between 1.14 and 1.47 seconds, and then plotted for each meridian with respect to eccentricity (Fig.5A). The standard deviations were slightly higher for the inferior meridian (±5.80nm inferior vs. ±8.13nm temporal). Two-tailed paired t-test revealed no significant difference between the two groups (p=0.71), indicating the similarity between the two meridians in the saturated amplitude of ΔOPL vs. time.

**Fig. 5.**
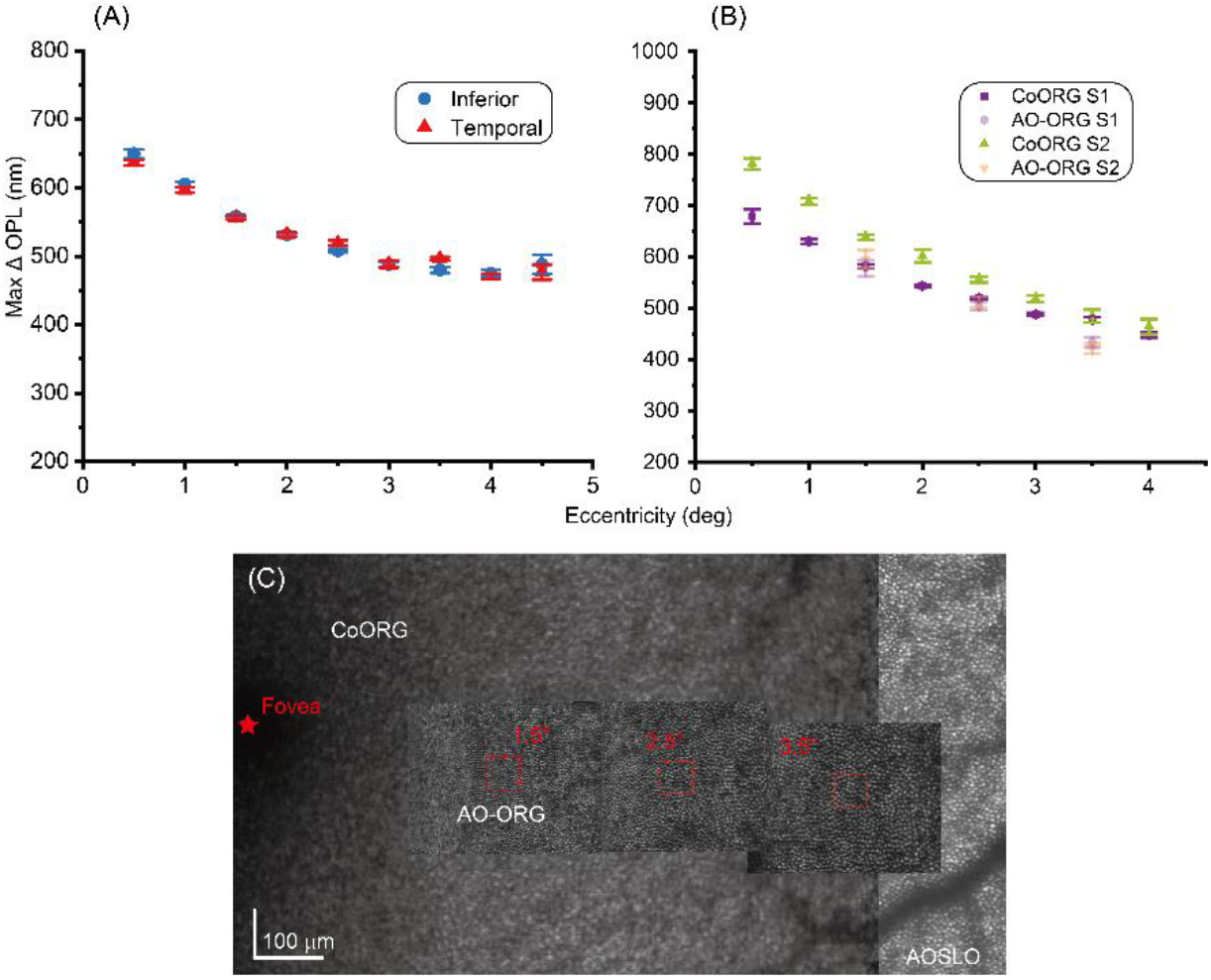
Comparison of OPL changes versus eccentricity for inferior and temporal meridians(A) and with AO and with CoORG (B) in the same subject. (C) *En face* overlaying image of CoORG, AO-ORG, and AOSLO, the size of red square is 0.3*0.3 deg^2^.

### 3.4 Comparison between CoORG and AO-based ORG

To compare the optical signals obtained from the extended field CoORG without AO against the AO-based ORGs obtained on a cellular scale, the saturated ΔOPL was compared between both at 1.5°, 2.5° and 3.5° inferior eccentricity following the same imaging protocol. The AO-OCT *en face* image at the ISOS was registered to the *en face* image of the same layer in CoORG (Fig.5C). A larger montage from an AOSLO (described previously[26]) aided with precise alignment between all modalities using blood vessel shadows as landmarks. Overall, ΔOPLs were similar between the AO-ORG and CoORG in the same subjects (Fig.5B). The averaged standard deviation across trials for the CoORG and AO-ORG are ±15.2nm and ±12.42nm for S1 and ±7.6mm and ±12.65mm for S2, respectively. There was high agreement between the ORGs at coarse-scale and those obtained at cellular resolution with AO. However, in S2, the AO based measurements were consistently lower in saturated ΔOPL with the maximum difference between the two being observed at 3.5° eccentricity equal to 62nm.

### 3.5 Inter individual variation in ORG

The mean saturated ΔOPL in the temporal meridian of 12 subjects is plotted in Figure 6, showing a decreasing trend with increasing eccentricity. The largest variation among subjects was noted at 0.5° eccentricity − ±88 nm and ± 66 nm (standard deviation)-for high and low bleach level, respectively. In general, the variability was lower with increasing eccentricity for the high bleach, but remained similar versus eccentricity for the low bleach. The range of standard deviation vs. eccentricity for high bleach was 88 to 31nm, and for low bleach was 66 to 17nm. One subject had a smaller dilated pupil size (6.82mm) and also a longer axial length (25.32mm). This led effectively to a lower photon density on the retina which is reflected in the significantly lower magnitude of ΔOPL in this subject (pink left triangle).

**Fig. 6.**
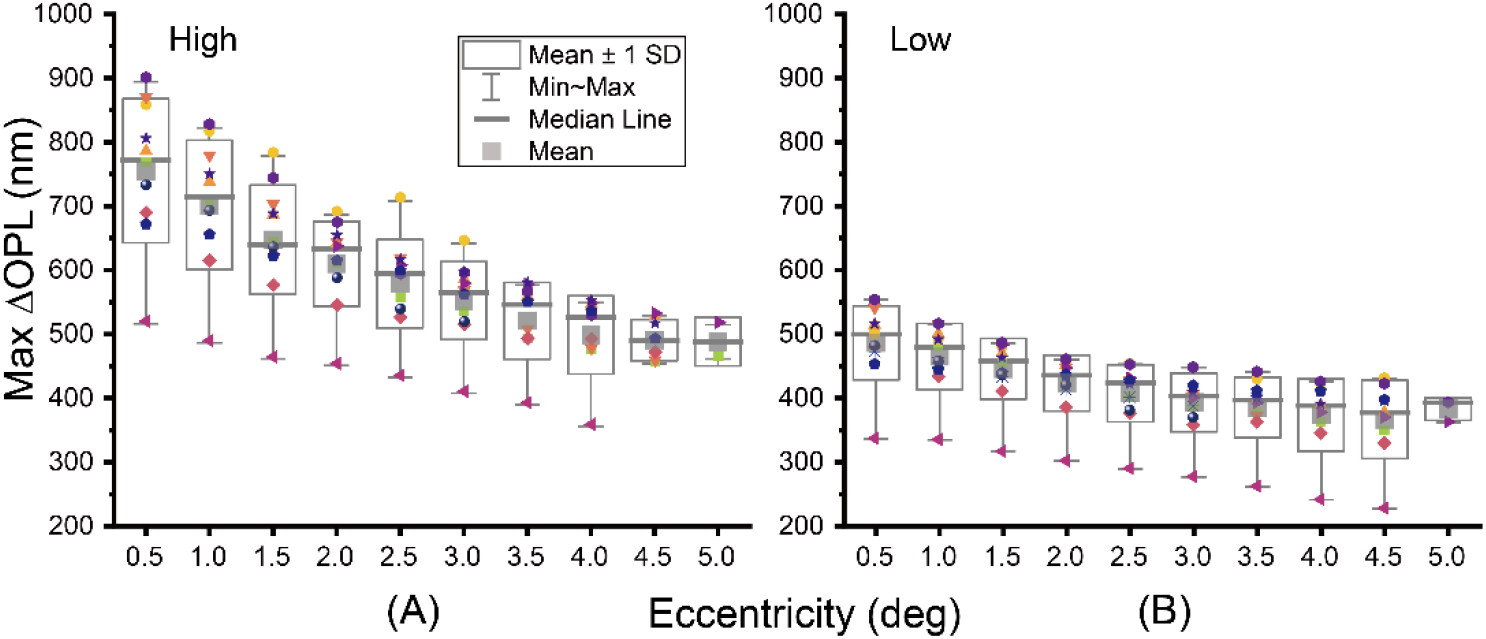
Box and whisker scatter plot with averaged maximum ΔOPL from 12 subjects for high and low levels. One subject‘s OPL was significantly lower than others (pink left triangle), due to a smaller pupil size and longer axial length translating effectively to a lower photon density on the retina.

Overall, the ORG varies among healthy subjects and we sought to assess the factors that contribute to this variation. A five-factor ANOVA was conducted to determine how five different variables affect ORG signal. Three of the factors were random factors (pupil size, axial length, and age; all characteristics of the subjects) while two of the factors were fixed factors (bleach level and eccentricity; specifically chosen for the experimental protocol), meaning that the expected mean squares were not standard and had to be calculated to determine the correct error term to use in each F-test [27]. This analysis revealed that age (p<0.001), bleach level (p<0.05), and eccentricity (p<0.001) individually were all significant factors in determining the saturated ΔOPL. The effect of age on ΔOPL, while statistically significant, seems to be driven by a larger variability in ORG response for the two younger age groups compared to a smaller variability in the older age group measurements, though this result may not be reproducible due to the small sample size for the older age group.

#### 3.5.1 Variation of ORG with photon density

Also of note, despite the two factors individually not significantly impacting ORG signal, the interaction between pupil size and axial length had a significant effect on ORG signal (p<0.001). To consider both pupil size and axial length factors together, photon density was calculated for each subject for each bleach level as mentioned in section 2.4. The saturated ΔOPL vs. eccentricity was divided across four bins of photon density, comprising both high and low bleach levels, and is plotted in Fig. 7. Paired t-test of two bins for high and low bleach level were calculated separately. Both were significantly different (p=0.001 for high bleach level and p=0.0008 for low bleach level), indicating that the saturated ΔOPL decreases with decreasing photon density. Closer to the fovea, the photon density has a larger impact on ΔOPL (range: 467 nm to 794 nm from lowest to highest photon density, a 1.7x change). With increasing eccentricity, this range reduced from 362 to 496 nm, going from the lowest to the highest photon density.

**Fig. 7.**
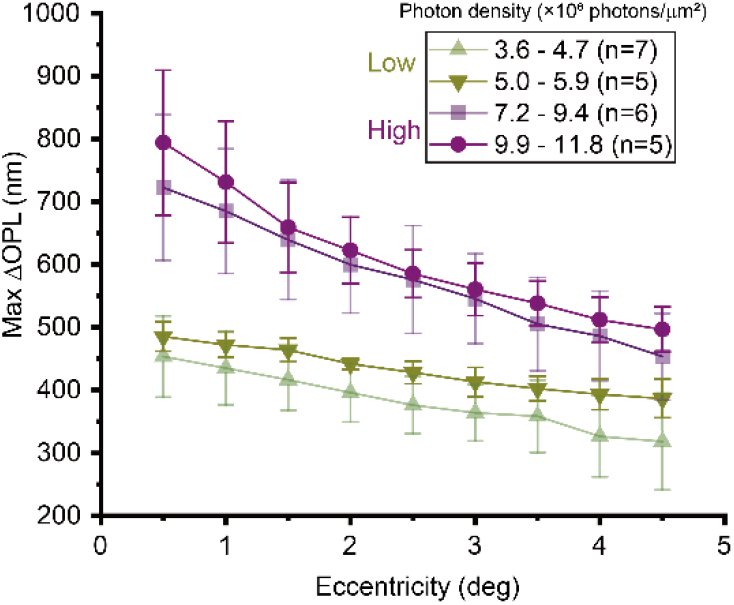
The variation in maximum saturated ΔOPL vs. eccentricity for different photon densities calculated specific to each subject. Data points are the mean in each bin, and error bars represent ± 1 standard deviation.

#### 3.5.2 Variation of ORG with eccentricity

The saturated ΔOPL at different bleach levels shows the same trend (seen in Figs. 5, 6 and 7), of decreasing amplitude with increasing eccentricity. Since these light evoked changes are contained within the cone outer segment, we sought to investigate whether variations in cone outer segment length versus eccentricity could explain this trend. The cone outer segment lengths were obtained from the CoORG B-scans for each subject. The saturated ΔOPL in 0.5 deg bins is plotted vs. the outer segment lengths for all subjects in Fig. 8. A linear regression applied to both high and low bleach conditions gave an R^2^ of 0.65 and 0.53 respectively. These coefficients of determination increased to 0.75 and 0.75 for the high and low bleach respectively removing the outlier who had significantly lower effective photon density on the retina (section 3.5). All regression coefficients were statistically significant (p<0.0001).

**Fig. 8.**
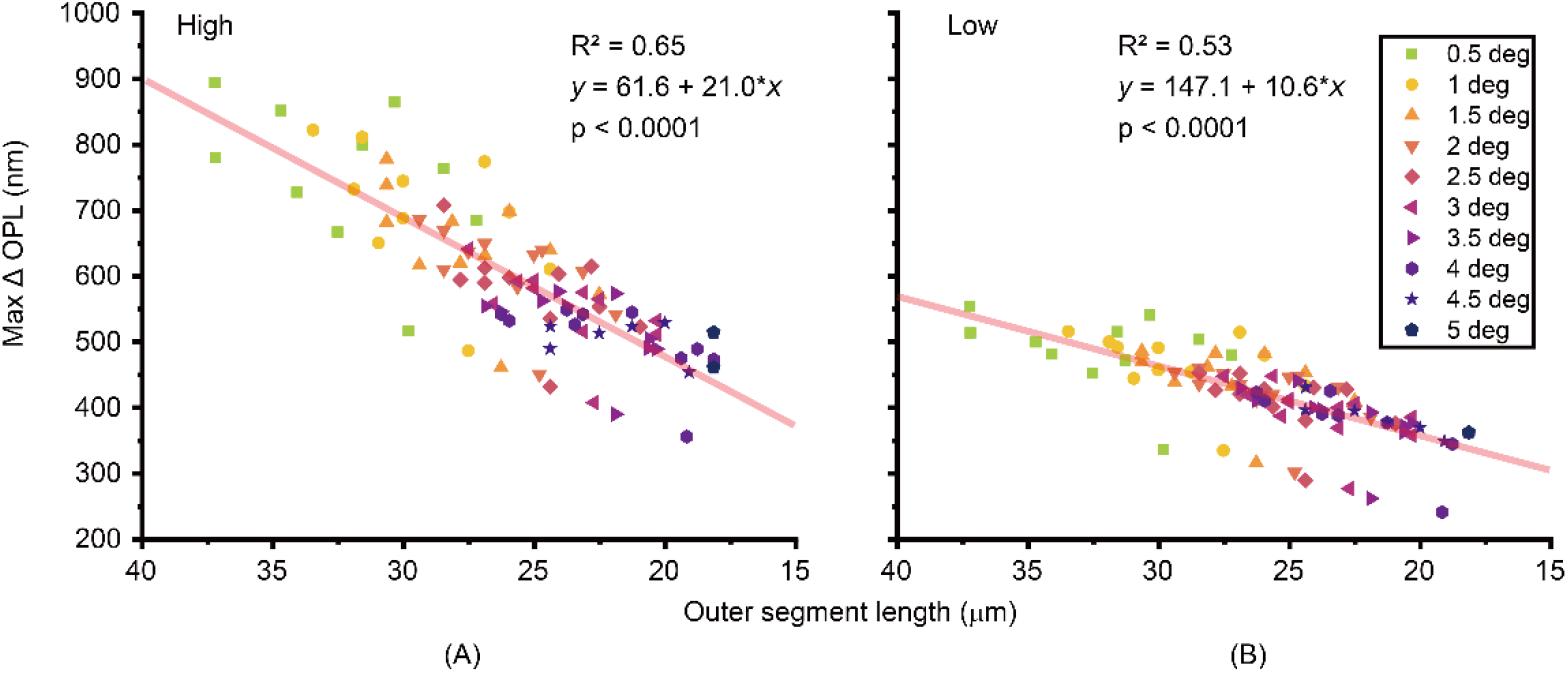
The variation in saturated ΔOPL versus outer segment length for high and low bleach strengths. Each data point indicates measurement from a single subject & eccentricity, while markers and colors indicate different eccentricity bins. A linear regression fit to all data is shown in both plots, along with the coefficient of determination and the equation for the linear fit.

## 4. Discussion

ORG has seen rapid development in the past few years, in a large part due to the enabling technologies of adaptive optics, OCT and SLO. Since it is non-invasive, and can assess retinal function with high sensitivity and spatial localization, there is high promise for this paradigm to replace the current gold standard of electroretinography (ERG). Given that a majority of ORG variants have employed AO, studies have been restricted to limited FOV and subjects. We introduce a coarse-scale ORG (CoORG) system without AO for extended FOV imaging and patient friendly operation. Given the high repeatability, rapid measurement, and good comparison with AO-based ORG measurements, the CoORG has high potential and applicability for clinical studies of age-related macular degeneration and inherited retinal disease. Using this paradigm, we created a normative cone ORG database of 12 subjects, one of the largest sized subject cohorts for ORG studies so far. Together, this not only facilitates comparative studies in diseased eyes using the same imaging platform, but also enables rapid assessment of how the inter-subject variability in anatomy or physiology impact the ORG. Overall, this brings us closer to the standardization of measurement and reporting protocols for ORG, similar in vein to the ISCEV standards established for ERGs[28].

To assess CoORG system performance, ORGs were recorded in 12 healthy subjects with two stimulus bleach levels. High trial-to-trial repeatability was observed across 5 repeat measurements. The general characteristics of the ORG closely followed published work using AO based ORG systems. To further compare the difference between CoORG and AO based ORG, the same subjects were imaged in both paradigms at different eccentricities indicating high degree of similarity between them

Furthermore, we characterized the potential impact of various factors affecting the ORG − retinal meridians, inter-subject variability, intra-subject variation vs. eccentricity and photon density, and cone outer segment length. The inferior and temporal meridians from the same subject showed no significant differences. Variations across subjects were observed due to differences in pupil size and axial length. ANOVA analysis suggested that while individually these two subject-specific factors do not have a significant impact, the interaction between them plays an important role in shaping the ORG response. This was expected since pupil area and axial length together govern the overall photon density on the retina. Previous work has detailed the positive correlation between the photon density and the magnitude & slope of ΔOPL[14]. This was confirmed here by accounting for each individual‘s pupil size and axial length in calculating the effective photon density. In the extreme case, the subject with the smallest pupil size and longest axial length, accordingly the lowest effective photon density on the retina, had a significantly smaller ΔOPL saturated amplitude for both bleach strengths. It is worth noting that a simplistic relationship based only on axial length was used to convert angular units to physical extent on the retina. In the future, a conversion including the anterior eye optical element surfaces - cornea and crystalline lens - may be used for a more precise estimate of photon density at the retina[29].

The variation of the ORG with eccentricity is noteworthy. The saturated ΔOPL[14] decreased with increasing eccentricity − a characteristic observed across all subjects and for a range of photon density (Fig 5,6, and 7). Photon density has a larger impact near the fovea compared to a larger eccentricity (Fig 7). These observations are explained to a large extent by the decreasing outer segment length versus eccentricity and examining its relationship with saturated ΔOPL. The saturated ΔOPL increases linearly with increasing outer segment length (Fig. 8). Importantly, the slopes of the fit suggest that the saturated ΔOPL magnitude is a small fraction of the outer segment length at all eccentricities − 0.02 % and 0.01% respectively for the high and low bleach. Therefore, the same photon density creates a larger absolute change in the fovea with the longer outer segments than that at a greater eccentricity. We chose to express the stimulus strength in photon density with units of photons per square micron as opposed to a bleach strength in percent or fraction. Key assumptions are made to convert photon density to bleach strength including waveguide condensation, pigment self-screening & optical density. It is unknown to what extent these assumptions vary across subjects. On the other hand, photon density provides a measure of stimulus strength largely independent of these assumptions and provides a favorable alternative, especially with a view towards standardizing the reporting protocols for ORG. The remaining factors that may potentially vary between subjects include absorption by the lens and macular pigment, for which, published values were used here. However, contribution of these factors at our stimulus center wavelength is low.

Some limitations remain to be improved in future work. The current stimulus beam size at the eye‘s pupil is larger than the average dilated pupil. The bright stimulus leads to pupil miosis preventing repeatable stimulus light dose at the retina and in turn negatively affecting the ORG repeatability. This issue was overcome here by cycloplegia. Pupil dilation can be avoided with a redesign of the stimulus assembly to focus the beam to a size smaller than 2-3 mm at the eye‘s pupil. To expand the FOV, lateral resolution was sacrificed. However, digital aberration correction, as has been implemented in full-field and line-scan OCT for instance, could be a powerful avenue for improving lateral resolution[30, 31].

Given the lack of cellular resolution, the CoORG cannot be a replacement for AO based ORG. The key advantage for AO in ORG lies in the ability to assess cellular scale function & dysfunction in photoreceptors and potentially other cell types in the future. This is important for both basic science and clinical applications. Since most vision restoration therapies function on a cellular scale, the resolution provided by AO is imperative for gauging their safety and efficacy. As a complement to the AO based ORG, the CoORG provides rapid, patient-friendly, high throughput functional imaging over an extended field. It can capture a 5° FOV ORG, with 5 repetitions to boost signal-to-noise in 10 minutes for a single bleach level. High speed operation without AO will enable such recordings in a range of pathological eyes with large eye movements, and in aging eyes where additionally the ocular media may be partially occluded by a cataract. In summary, the CoORG provides a powerful and complementary avenue to the AO based ORG for translating functional imaging to the clinic.

## Data availability

Data underlying the results presented in this paper are not publicly available at this time but maybe obtained from the authors upon reasonable request.

## References

1. Mulligan, J., The Optoretinogram at 38. Journal of Vision, 2019. 19(8): p. 33–33.

2. Grieve, K. and A. Roorda, Intrinsic signals from human cone photoreceptors. Investigative ophthalmology & visual science, 2008. 49(2): p. 713–719.

3. Cooper, R.F., et al., Non-invasive assessment of human cone photoreceptor function. Biomedical optics express, 2017. 8(11): p. 5098–5112.

4. Cooper, R.F., D.H. Brainard, and J.I.W. Morgan, Optoretinography of individual human cone photoreceptors. Optics Express, 2020. 28(26): p. 39326–39339.

5. Jonnal, R.S., et al., In vivo functional imaging of human cone photoreceptors. Optics express, 2007. 15(24): p. 16141–16160.

6. Rha, J., et al., Variable optical activation of human cone photoreceptors visualized using a short coherence light source. Optics letters, 2009. 34(24): p. 3782–3784.

7. Kim, T.-H., et al., Functional optical coherence tomography enables in vivo optoretinography of photoreceptor dysfunction due to retinal degeneration. Biomedical Optics Express, 2020. 11(9): p. 5306–5320.

8. Yao, X.-C., et al., Rapid optical coherence tomography and recording functional scattering changes from activated frog retina. Applied optics, 2005. 44(11): p. 2019–2023.

9. Bizheva, K., et al., Optophysiology: depth-resolved probing of retinal physiology with functional ultrahigh-resolution optical coherence tomography. Proceedings of the national academy of sciences, 2006. 103(13): p. 5066–5071.

10. Srinivasan, V., et al., In vivo measurement of retinal physiology with high-speed ultrahigh-resolution optical coherence tomography. Optics letters, 2006. 31(15): p. 2308–2310.

11. Zhang, P., et al., In vivo optophysiology reveals that G-protein activation triggers osmotic swelling and increased light scattering of rod photoreceptors. Proceedings of the National Academy of Sciences, 2017. 114(14): p. E2937–E2946.

12. Hillmann, D., et al., In vivo optical imaging of physiological responses to photostimulation in human photoreceptors. Proceedings of the National Academy of Sciences, 2016. 113(46): p. 13138–13143.

13. Pandiyan, V.P., et al., High-speed adaptive optics line-scan OCT for cellular-resolution optoretinography. Biomedical Optics Express, 2020. 11(9): p. 5274–5296.

14. Pandiyan, V.P., et al., The optoretinogram reveals the primary steps of phototransduction in the living human eye. Science Advances, 2020. 6(37): p. eabc1124.

15. Pandiyan, V.P., et al., Reflective mirror-based line-scan adaptive optics OCT for imaging retinal structure and function. Biomedical optics express, 2021. 12(9): p. 5865–5880.

16. Jonnal, R.S., et al., Phase-sensitive imaging of the outer retina using optical coherence tomography and adaptive optics. Biomedical optics express, 2012. 3(1): p. 104–124.

17. Zhang, F., et al., Cone photoreceptor classification in the living human eye from photostimulation-induced phase dynamics. Proceedings of the National Academy of Sciences, 2019. 116(16): p. 7951–7956.

18. Azimipour, M., et al., Optoretinogram: optical measurement of human cone and rod photoreceptor responses to light. Optics letters, 2020. 45(17): p. 4658–4661.

19. Pfäffle, C., et al., Simultaneous functional imaging of neuronal and photoreceptor layers in living human retina. Optics Letters, 2019. 44(23): p. 5671–5674.

20. Pfäffle, C., et al. Phase-sensitive measurements of depth dependent signal transduction in the inner plexiform layer. in Ophthalmic Technologies XXXI. 2021. SPIE.

21. Lassoued, A., et al., Cone photoreceptor dysfunction in retinitis pigmentosa revealed by optoretinography. Proceedings of the National Academy of Sciences, 2021. 118(47): p. e2107444118.

22. Teng, P.-y., Caserel-an open source software for computer-aided segmentation of retinal layers in optical coherence tomography images. Zenodo, DOI, 2013. 10.

23. Stevenson, S.B. and A. Roorda. Correcting for miniature eye movements in high-resolution scanning laser ophthalmoscopy. in Ophthalmic Technologies XV. 2005. International Society for Optics and Photonics.

24. Norren, D.V. and J.J. Vos, Spectral transmission of the human ocular media. Vision research, 1974. 14(11): p. 1237–1244.

25. Stockman, A., L.T. Sharpe, and C. Fach, The spectral sensitivity of the human short-wavelength sensitive cones derived from thresholds and color matches. Vision research, 1999. 39(17): p. 2901–2927.

26. Jiang, X., et al., Measuring and compensating for ocular longitudinal chromatic aberration. Optica, 2019. 6(8): p. 981–990.

27. Kirk, R.E., Experimental design: Procedures for the behavioral sciences. 2012: Sage Publications.

28. McCulloch, D.L., et al., ISCEV Standard for full-field clinical electroretinography (2015 update). Documenta ophthalmologica, 2015. 130(1): p. 1–12.

29. Huang, X., T. Anderson, and A. Dubra, Retinal magnification factors at the fixation locus derived from schematic eyes with four individualized surfaces. Biomedical Optics Express, 2022. 13(7): p. 3786–3808.

30. Hillmann, D., et al., Aberration-free volumetric high-speed imaging of in vivo retina. Scientific reports, 2016. 6(1): p. 1–11.

31. Ginner, L., et al., Noniterative digital aberration correction for cellular resolution retinal optical coherence tomography in vivo. Optica, 2017. 4(8): p. 924–931.

